# Transgenerational inheritance of abnormal spermatogenesis with *Igf2/H19* epigenetic alteration in CD1 mouse induced by *in utero* arsenic exposure

**DOI:** 10.1101/2020.04.16.044032

**Authors:** Guoying Yin, Liting Xia, Yaxing Hou, Yaoyan Li, Deqing Cao, Yanan Liu, Jingshan Chen, Juan Liu, Liwen Zhang, Qiaoyun Yang, Qiang Zhang, Naijun Tang

**Author notes:** Corresponding author: Qiang Zhang, Department of Occupational and Environmental Health, School of Public Health, Tianjin Medical University, No. 22 Qixiangtai Road, Heping District, Tianjin, 300070, China.

## Abstract

Developmental exposure to environmental toxicants can induce transgenerational reproductive disease phenotypes through epigenetic mechanisms. However, little is known about the transgenerational effects of arsenic exposure. We hypothesize that prenatal arsenic exposure may result in impaired spermatogenesis in subsequent generations of male mice. To test our hypothesis, we treated pregnant CD-1 (F0) mice with drinking water containing sodium arsenite (85 ppm) from days 8 to 18 of gestation. Male offspring were bred with untreated female mice until the F3 generation was produced. Our results revealed that transient exposure of the F0 gestating female to arsenic can result in decreased sperm quality and histological abnormalities in testes of male offspring in the F1 and F3 generations. The overall methylation status of *Igf2* DMR2 and *H19* DMR was significantly lower in the arsenic-exposed group than that of the control group in both F1 and F3 generations. The relative mRNA expression levels of *Igf2* and *H19* in arsenic-exposed males were significantly higher than in the control males in both F1 and F3 generations. This study indicates that ancestral exposure to arsenic may result in transgenerational inheritance of an impaired spermatogenesis phenotyping involving both epigenetic alterations and the abnormal expression of *Igf2* and *H19*.

## 1. Introduction

Arsenic is a natural element of the Earth’s crust and is widely distributed in the environment as a result of both natural sources and human activities. It is reported that hundreds of millions of people suffer from arsenic poisoning from drinking water worldwide (Murcott 2012). Numerous studies have shown that chronic exposure to inorganic arsenic (iAs) is associated with various cancers including skin, lung, liver, and bladder cancers (Bustaffa et al. 2014), as well as other non-cancer disorders including cardiovascular diseases, metabolic diseases, and mental disorders (Tolins et al. 2014; Kuo et al. 2017).

Arsenic is an endocrine disruptor and many studies have suggested arsenic as a reproductive and developmental toxicant. In various experimental models, arsenic-induced male reproductive toxicity is evidenced by a decrease in testes weight and diminished sperm count and motility, as well as increased sperm malformation (Jana et al. 2006; Chang et al. 2007; Sanghamitra et al. 2008). Several studies have reported that arsenic mainly impairs spermatogenesis via aberrant modulation of male reproductive hormones, especially testosterone (Alamdar et al. 2017). In humans, arsenic exposure is associated with low semen quality (Oguri et al. 2016), infertility (Wang et al. 2016), and erectile dysfunction in men (Hsieh et al. 2008). There is increasing evidence suggesting that *in utero* arsenic exposure poses a serious health risk to the developing fetus and newborn (Quansah et al. 2015; Shih et al. 2017). Moreover, early life arsenic exposure increases the risk of disease development in offspring during childhood and later in life (Farzan et al. 2013). Mouse models have been used to study the carcinogenic effects of arsenic exposure *in utero* at high levels (42.5 and 85 ppm) (Waalkes et al. 2007; Tokar et al. 2012). Using the same experimental model, Nohara et al. (2017) found that *in utero* arsenic exposure increased hepatic tumors in the F1 and F2 male offspring. Although this paradigm has been used to study arsenic-related obesity in female offspring of CD-1 mice (Rodriguez et al. 2016), the reproductive effects of *in utero* arsenic exposure on subsequent generations remain unclear. Therefore, this study aims to explore the potential transgenerational reproductive toxicity of male mice exposed to high levels of arsenic *in utero*.

Transgenerational epigenetic inheritance is defined as germline-mediated transmission of environmentally induced phenotypes between generations in the absence of direct environmental exposure. When a gestating female (F0 generation) is exposed to an environmental compound, the F1 generation embryo and F2 generation germline were directly exposed to the environmental compound, whereas the F3 generation was the first generation not directly exposed (Heard and Martienssen 2014; Martos et al. 2015). Recent studies have shown that exposure to endocrine-disrupting chemicals can alter epigenetic marks in the germline and induce male reproductive toxicity of multiple generations (Skinner 2016). More recently, it has been reported that maternal arsenite exposure may induce transgenerational reproductive toxicity in *Caenorhabditis elegans* by demethylation of histone H3K4 (Yu and Liao 2016). However, the transgenerational male reproductive effect of prenatal arsenic exposure remains to be studied in mammals.

Imprinted genes have been shown to correlate with transgenerational epigenetic inheritance (Stouder and Paoloni-Giacobino 2010). They are often found in clusters and differentially methylated regions (DMRs) are key for imprinted gene expression (Reik and Walter 2001). In mammals, the period of DNA methylation reprogramming (erasure and reestablishment) is a critical window of sensitivity to environmental exposures. Epigenetic markers induced by environmental toxicants during primordial germ cell (PGC) differentiation can be transgenerationally perpetuated (Bohacek and Mansuy 2015). Recent studies have shown that a loss of methylation in the insulin-like growth factor 2 (*Igf2*) DMR2 or in both the *Igf2* DMR2 and the *H19* DMR was associated with quantitative defects of spermatogenesis (Boissonnas et al. 2010). Furthermore, *in utero* arsenic exposure may adversely influence DNA methylation profiles at various cytosine phosphate guanines (CpGs) (Kaushal et al. 2017).

In this study, we hypothesized that a transient exposure to arsenic could induce impaired spermatogenesis by altering epigenetic profiles of imprinted genes. We used an outbred mouse model and a transgenerational scheme originating from *in utero* exposure to high-dose sodium arsenite (85 ppm). To reveal the possible transgenerational phenotype and underlying mechanism, we evaluated the sperm quality and spermatogenesis of the male offspring and quantified the methylation pattern of the specific imprinted genes *Igf2* DMR2 and *H19* DMR and their relative expression in the testis of F1 and F3 generations.

## 2 Materials and standards

### 2.1 Animals and treatment

All experimental protocols for the procedures with mice were preapproved by the Ethics Committee of Tianjin Medical University. The animals were treated humanely and all efforts were made to avoid their suffering.

Seven-week-old male and female CD-1 mice were purchased from Beijing Vital River Laboratory Animal Technology Co., Ltd (Beijing, China) and housed in a specific pathogen-free environment under conditions of controlled temperature (22 ± 2°C), humidity (50% to 60%), and a 12:12-h light:dark cycle. After 1 week of acclimation, female mice were timed-mated with male mice at a ratio of 2:1. The date of vaginal plug detection was designated as gestation day 0 (GD 0). The pregnant female mice (F0 generation) were assigned randomly to one of two groups (six pregnant females per group): a) control group (control lineages), without arsenic exposure; b) arsenic-exposed group (As lineages), with 85 ppm sodium arsenite obtained from Sigma-Aldrich (St Louis, MO, USA). Arsenic was administered in the drinking water and the solution was prepared daily to avoid oxidation. The treatment window was continued from GD8 to GD18. Daily water and food consumption were recorded from GD0 to GD18. The weights of the pregnant mice were recorded at GD0 and GD18, respectively. After spontaneous parturition, the F1 offspring were fostered with dams throughout lactation. At weaning, the F1 male mice were separated from their female littermates, individually housed, and given food and water *ad libitum*. Eight-week-old male mice from the F1 generation of control or As lineages were timed-mated with wild-type female mice to obtain F2 generation offspring. The F2 generation male mice were timed-mated with wild-type female mice to obtain F3 generation offspring.

Adult male mice at 13 weeks of age (around postnatal day 90) were sacrificed for tissue harvest. Body and organ weights were measured at dissection time. For histopathology examination, the left testis was removed, weighed, and fixed in 4% paraformaldehyde. For gene and protein analysis, the right testis was quickly removed, weighed, and flash-frozen in liquid nitrogen, then stored at −80°C until use.

### 2.2 Sperm quality analysis

Sperm quality was determined by motility and malformation. An equal length of the left cauda epididymides was separated and minced with scissors in an EP tube containing 1 mL of 0.9% saline. The tube was placed in a 37°C water bath for 30 min to allow the sperm to be released from the cut epididymis into saline sufficiently. The sperm suspension was pipetted into a pre-warmed (37°C) hemocytometer. Then, sperm motility was analyzed by counting the motile and immotile spermatozoa under the light microscope and expressed as percentage of the total sperm counts. All analysis was performed by the same trained observer to avoid subjective differences in motility evaluation. For sperm malformation analysis, the suspension was smeared onto clean slides and allowed to dry. Slides were then fixed in methanol for 10 min at room temperature and air-dried. The film preparations were stained with hematoxylin and eosin and the ratio of sperm malformation was calculated by counting the number of malformed spermatozoa (including head, midpiece and tail defects) per 1000 sperm in each sample under light microscope. Six animals from different lineages were used for sperm quality analysis in each group of F1 and F3 generations.

### 2.3 Histopathological analysis of testis and quantitative evaluation of spermatogenesis

The fixed testis was trimmed, dehydrated, and embedded in paraffin blocks by standard procedures. Tissue sections of 5-μm thickness were made and stained with hematoxylin and eosin for histopathology analysis under an optical microscope equipped with a digital camera (IX81, Olympus, Japan). Spermatogenesis is routinely divided into 12 stages (I-XII stages) in mice, which is characterized by the shape of acrosomes according to known criteria (Meistrich and Hess 2013). In this study, quantitative evaluation of spermatogenesis was carried out by counting the relative number of each type of germ cells at stage VII of the seminiferous cycle. The types of germ cells include type A spermatogonia (Asg), preleptotine spermatocytes (pLSc), mid-pachytene spermatocytes (mPSc) and step 7 spermatids (7Sd). Briefly, for each section of the mouse testis, all germ cells present in stage VII of the cycle were counted in 10 round or nearly round seminiferous tubules. In addition, maturity of the germinal epithelium was evaluated by Johnsen score (Johnsen 1970), a simple way for assessment of spermatogenesis. For the Johnsen score a grade from 1 to 10 was given to each tubule cross-section according to the range from no cells to complete spermatogenesis (**Table S1**). For each animal, 50 tubules were evaluated. Six animals per group of F1 and F3 generations were analyzed.

### 2.4 Bisulfite sequencing PCR for *H19* DMR and *Igf2* DMR2

Bisulfite treatment of the DNA of testis was done according to the instructions of the EZ DNA Methylation-Direct™ Kit (Zymo Research, USA), which features simple and reliable DNA bisulfite conversion directly from tissue without the prerequisite for DNA purification. Nested PCR using ZymoTaq™ qPCR PreMix (Zymo Research) was performed for the amplification of the *Igf2* DMR2 and *H19* DMR. The primer sequences are listed in **Table 1**. The PCR products were recovered and gel-purified using Zymoclean Gel DNA Recovery Kit (Zymo Research). The purified DNA fragments were ligated into the pBLUE-T vector and transformed into *Escherichia coli* according to the manufacturer’s instructions. After incubation, 10 to 20 positive clones were selected from each sample and sequenced on ABI Prism 310 (Applied Biosystems, USA). Ten effective DNA methylation sequencing data per sample were randomly selected and analyzed by BiQ analyzer.

**Table 1.**
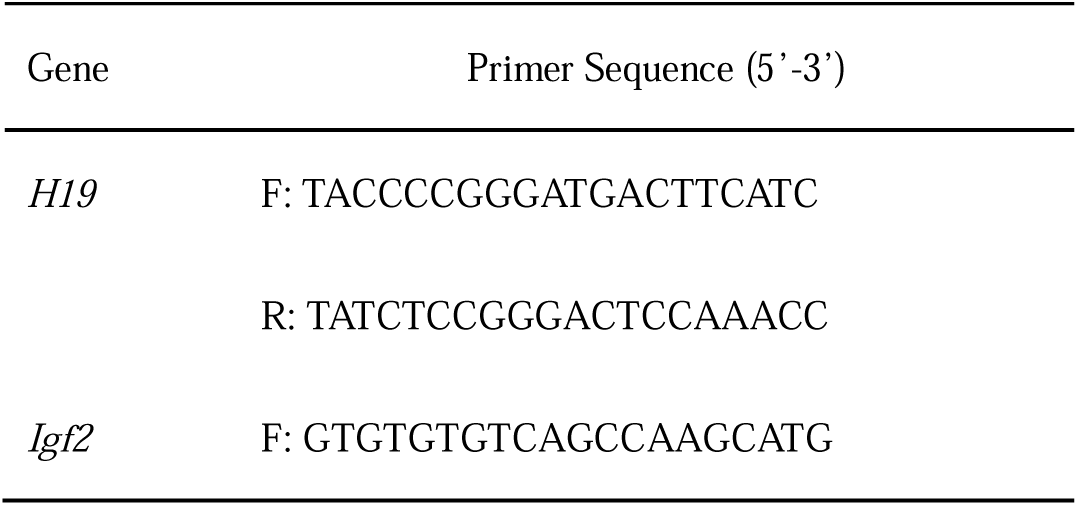
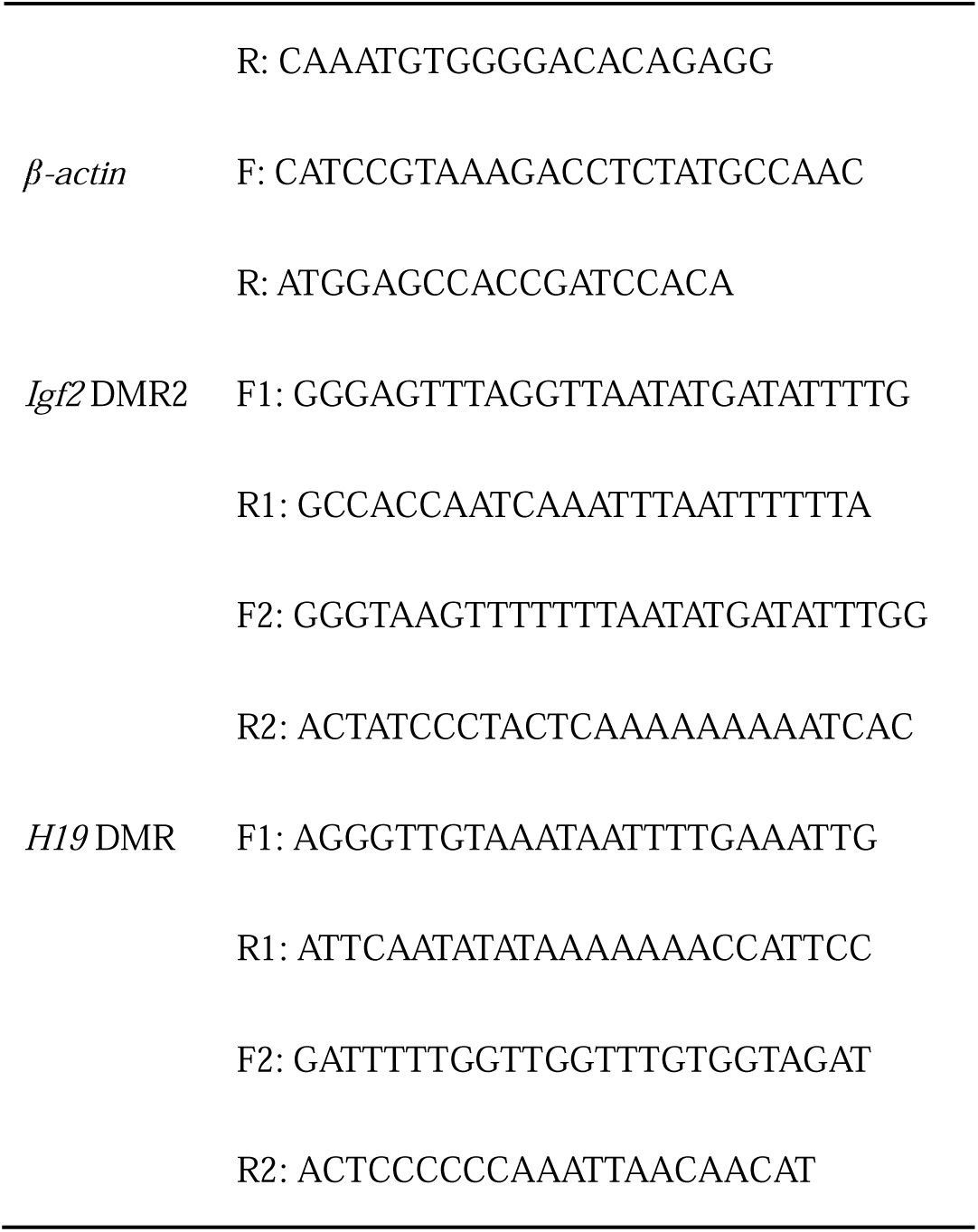
Primers used for qRT-PCR and bisulfte sequencing.

### 2.5 Quantitative real-time polymerase chain reaction analysis of *Igf2* and *H19*

Total RNA was extracted from testis tissues by using TRIzol® Reagent (Invitrogen, USA) according to the manufacturer’s instructions. cDNA was synthesized by reverse transcription of 0.5 µg RNA using PrimeScript RT Master Mix (Takara, Japan). qRT-PCR was performed in a 20 μL reaction system containing 10 μL 2 × SYBR® Premix Ex Taq II (Takara, Japan), 0.8 μL of the polymerase chain reaction (PCR) forward primer (10 μmol/L), 0.8 μL of the PCR reverse primer (10 μmol/L), 2 μL of the cDNA sample, and 6.4 μL dH_2_O. The amplification was performed using LightCycler®480 Instrument (Roche, Switzerland). The reaction conditions are as follows: 1 cycle at 95°C for 30 s; followed by 40 cycles at 95°C for 5 s and 60°C for 30 s; and then 1 cycle at 95°C for 5 s, 60°C for 60 s and 95°C; and finally, 1 cycle at 50°C for 30 s. The specificity of the PCR product was verified by melting curves. *β*-actin was used as a housekeeping gene for quantitative analysis. The expression levels of individual target genes were quantified with 2–ΔΔCt method. All experiments on gene expression were performed in triplicate. Primer sequences are shown in **Table 1**.

### 2.6 Statistical analysis

The SPSS software (version 16.0, Chicago, IL, USA) was used for the statistical analysis. The quantitative data were analyzed by Student *t*-test and the qualitative data were analyzed by chi-square test or Fisher’s exact test. The values are presented as mean ± SD or percentages (%). A statistically significant difference was defined as *p* < 0.05.

## 3. Results

### 3.1 F0 maternal water and food intake as well as body weight gain

The mean weights of F0 pregnant female mice at GD 0 and GD 18 were not statistically different between control and arsenic-exposed groups. However, the difference in mean weight gain (33.1 g control; 28.2 g As) was statistically different (**Fig. 1A**). The average water consumption for the control mice was 7.2 mL/day before GD8 and 9.1 mL/day after GD8. The water consumption for the arsenic treated mice decreased from 7.3 mL/day before GD8 to 4.7 mL/day after GD8. The average daily water consumption of the arsenic-exposed group was significantly lower than that of the control group during the entire gestation period (**Fig. 1B**). The daily food consumption was similar between the two groups (**Fig. 1C**).

**Fig. 1.**
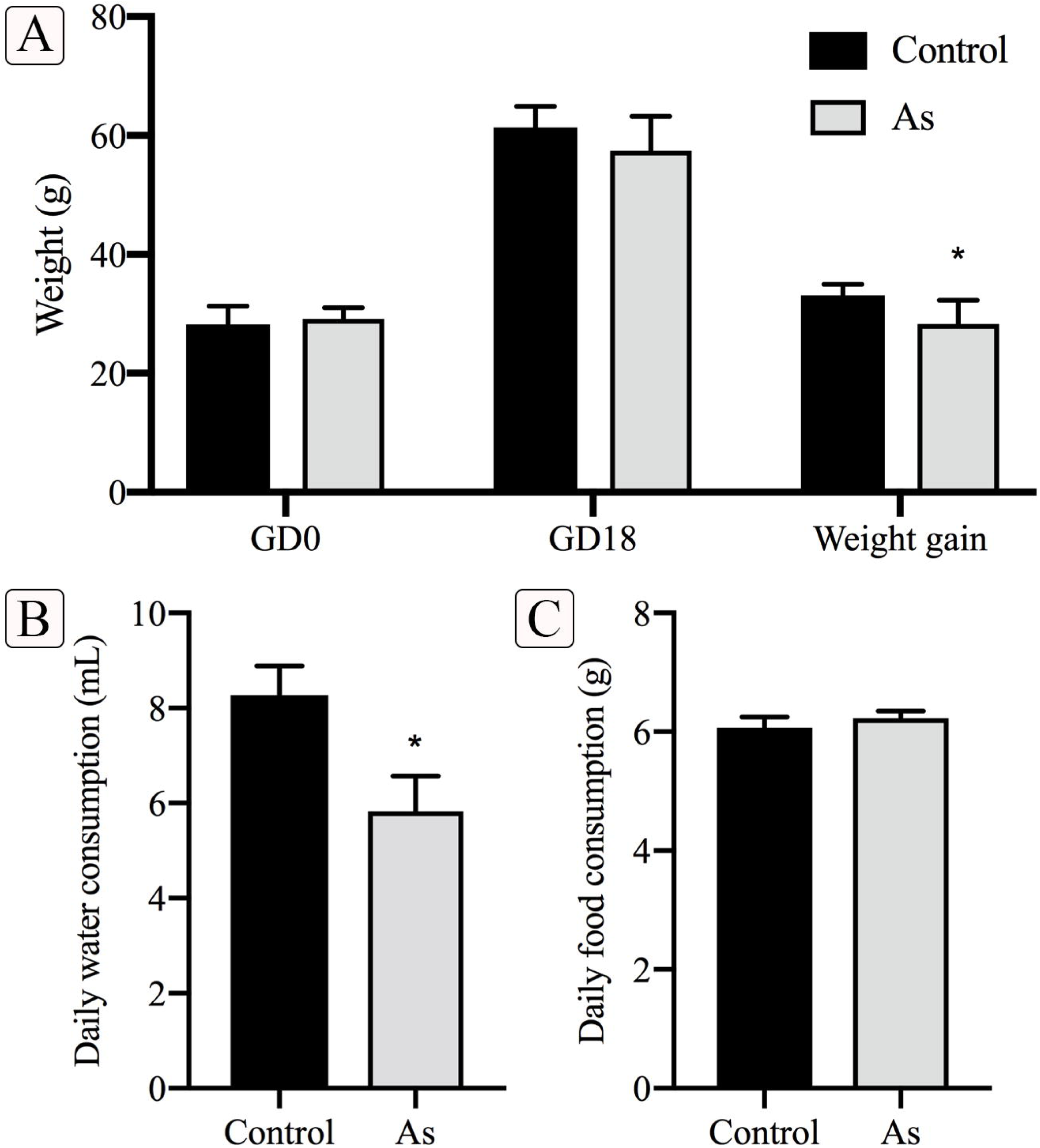
The impact of *in utero* arsenic exposure on maternal food and water intake as well as body weight gain in pregnant CD-1 mice. **A** Body weight of F0 pregnant mice at GD0 and GD18, as well as body weight gain during the pregnancy (from GD0 to GD18). **B** Daily water consumption of F0 pregnant mice during the pregnancy. **C** Daily food consumption of F0 pregnant mice during the pregnancy. Each bar represents the mean ± SEM (*n* = 6). * *p* < 0.05

### 3.2 Sperm quality of F1 and F3 male offspring

We used sperm quality to determine if there were male reproductive effects on future generations. In the F1 generation, the sperm motility of the arsenic-exposed group (81.2 ± 8.6%) was significantly lower than that in the control group (91.1 ± 3.2%; *p* < 0.05) (**Fig. 2A**). For the F3 generation, the sperm motility of the arsenic-exposed group was still lower than that of the control group, but the difference was not statistically significant. In contrast, the sperm malformation rate of the arsenic-exposed group was significantly higher than that of the control group in both F1 and F3 generations (*p* < 0.05) (**Fig. 2B**). We did not detect any differences in the litter size or the female/male ratio in the offspring **(Table S2)**.

**Fig. 2.**
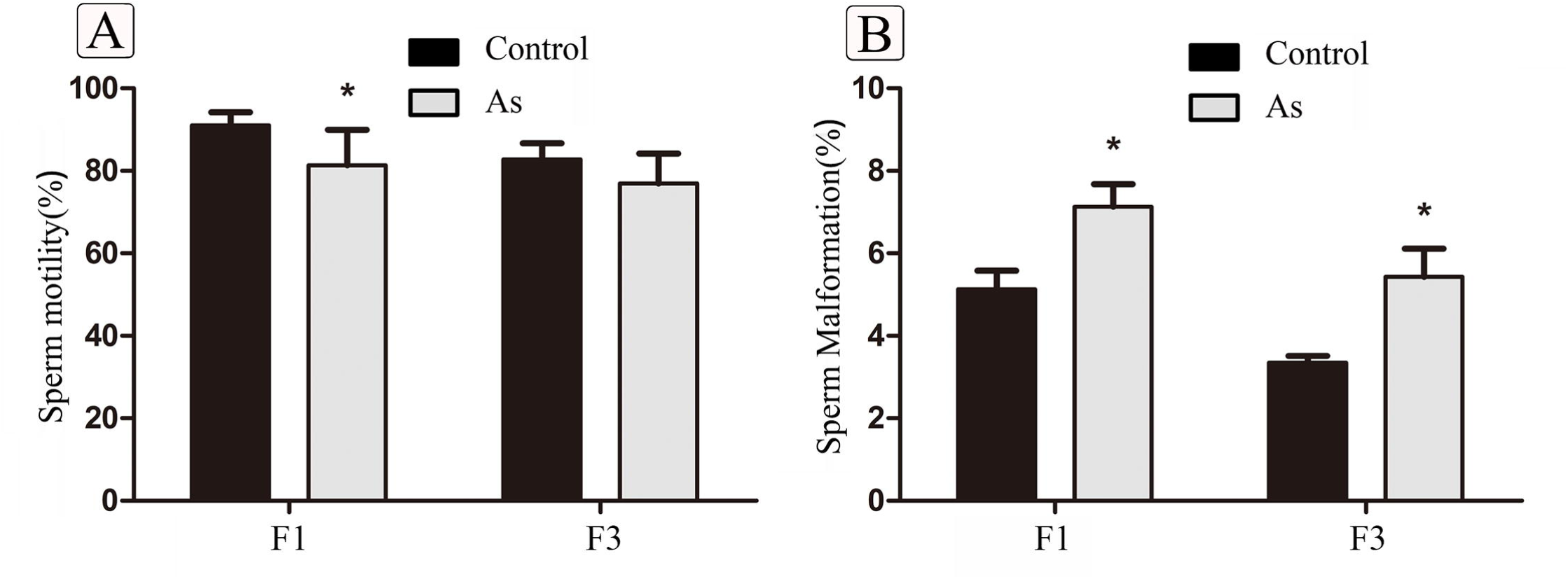
Transgenerational effects of *in utero* arsenic exposure on male sperm motility and malformation. **A** The sperm motility of the arsenic-exposed and control groups in F1 and F3 generations. **B** The sperm malformation rate of the arsenic-exposed and control groups in F1 and F3 generations. Each bar represents the mean ± SEM (*n* = 6). * *p* < 0.05

### 3.3 Histological changes in testis of F1 and F3 male offspring

The testes weight and epididymides weight were similar between the two groups in both F1 and F3 generations. However, the difference in testes/body weight ratio between the arsenic-exposed group and the control group was statistically significant in the F1 generation (**Table S3**). Testicular sections of the control mice showed normal cell arrangement in the seminiferous tubules including developing germ cells at different stages (spermatogonia, spermatocytes, spermatids, and matured spermatids) in the stratified epithelium and in the lumen of tubules **(Fig. 3A and a, C and c)**. However, significant histological changes appeared in the testes of arsenic-exposed F1 male mice, such as degeneration of seminiferous tubules, reduced layers of germ cells, intercellular dissociation of spermatogenic cell lines, and decreased number of spermatozoa **(Fig. 3B and b)**. Although the histological changes of arsenic-exposed F3 mice were not as obvious, the layers and number of germ cells were reduced **(Fig. 3D and d)**.

**Fig. 3.**
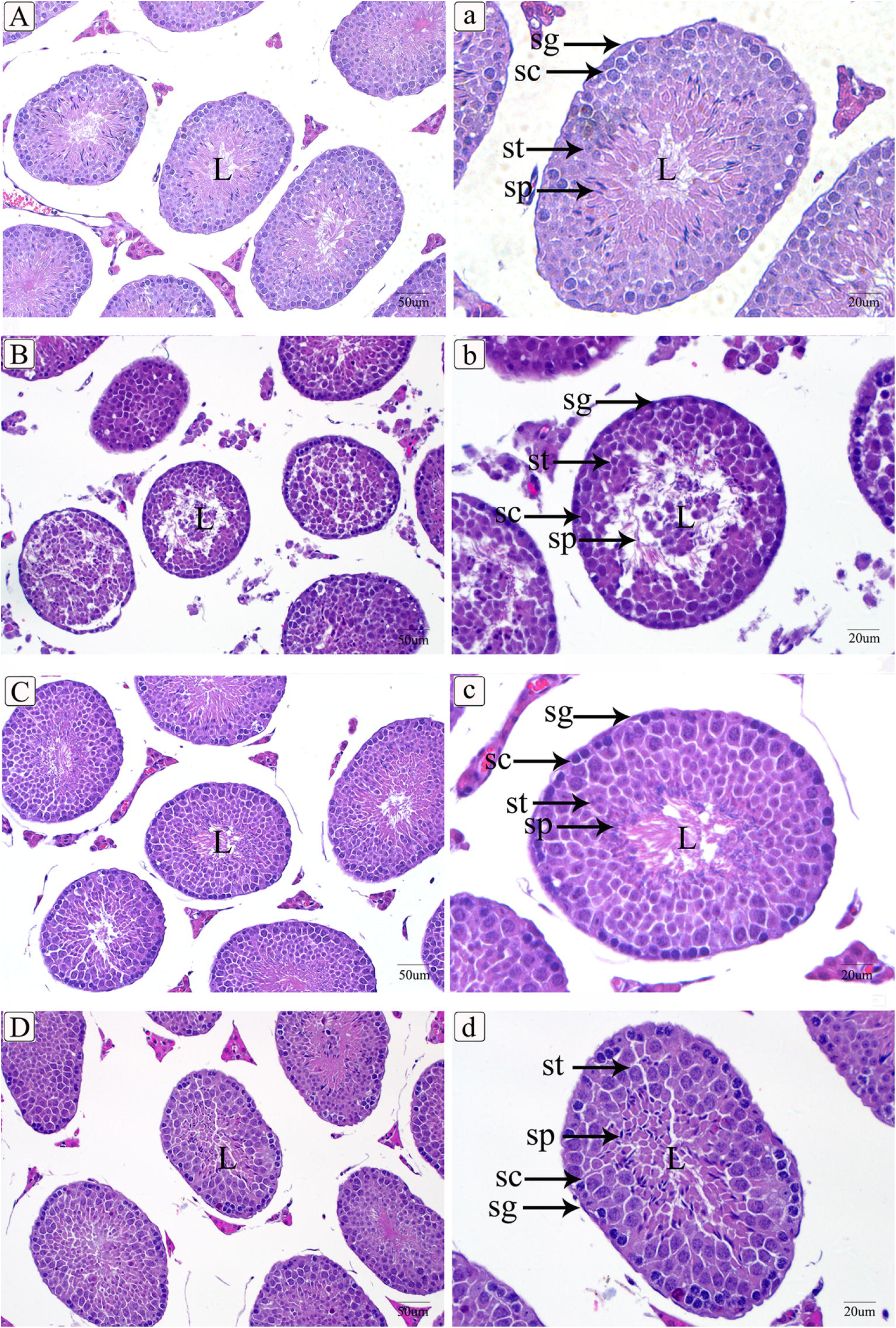
Photomicrographs of seminiferous tubules including developing germ cells at different stages (spermatogonia, spermatocytes, spermatids, and matured spermatids) in the stratified epithelium and in lumen of tubules. A and a control groups in F1 generation, B and b arsenic-exposed groups in F1 generation, C and c control groups in F3 generation, D and d arsenic-exposed groups in F3 generation. H&E stain. A, B, C, D scale bar 50 μm; a, b, c, d scale bar 20 μm.

### 3.4 Abnormal spermatogenesis in testis of F1 and F3 male offspring

The quality of the seminiferous epithelium was assessed by the Johnsen scoring system. For F1 generation, Johnsen score in the control group was 9.0 ± 0.6 and for the arsenic-exposed group, it was 7.6 ± 0.3. For the F3 generation, it was 9.0 ± 0.5 and 8.1 ± 0.2 for the control and arsenic-exposed groups, respectively. Johnsen score showed significant reduction in the arsenic-exposed group in both F1 and F3 generations, which represented poor spermatogenesis (*p* < 0.05) (**Fig. 4**). We further examined spermatogenesis by counting the relative number of different types of germ cells at stage VII of the spermatogenic cycle. The number of Asg, mPSc, and 7Sd was decreased in the arsenic-exposed group compared with that of the control group in both F1 and F3 generations. Notably, a significant decrease was shown in the number of mPSc in F1 generation and Asg in F3 generation of the arsenic-exposed group compared with the control group (**Table 2**). These findings indicate that ancestral exposure to arsenic may result in transgenerational inheritance of an increased susceptibility to disturbed spermatogenesis of male CD-1 mice in subsequent generations.

**Table 2.** The number of Asg, pLSc, mPSc, and 7Sd at stage VII of the spermatogenic cycle (n = 6, mean ± SD)

| Generation | Group | Asg | pLSc | mPSc | 7Sd |
| --- | --- | --- | --- | --- | --- |
| F1 | Control | $6.73 \pm 1.97$ | $32.96 \pm 7.02$ | $60.26 \pm 8.12$ | $140.63 \pm 21.00$ |
| | As | $5.92 \pm 1.37$ | $34.64 \pm 8.42$ | $50.98 \pm 8.07^*$ | $133.94 \pm 20.53$ |
| F3 | Control | $7.37 \pm 1.85$ | $35.41 \pm 6.05$ | $50.44 \pm 5.21$ | $136.25 \pm 10.12$ |
| | As | $5.05 \pm 0.55^*$ | $30.74 \pm 4.38$ | $48.55 \pm 5.35$ | $125.45 \pm 12.61$ |

\* Compared with the control group: $p < 0.05$

**Fig. 4.**
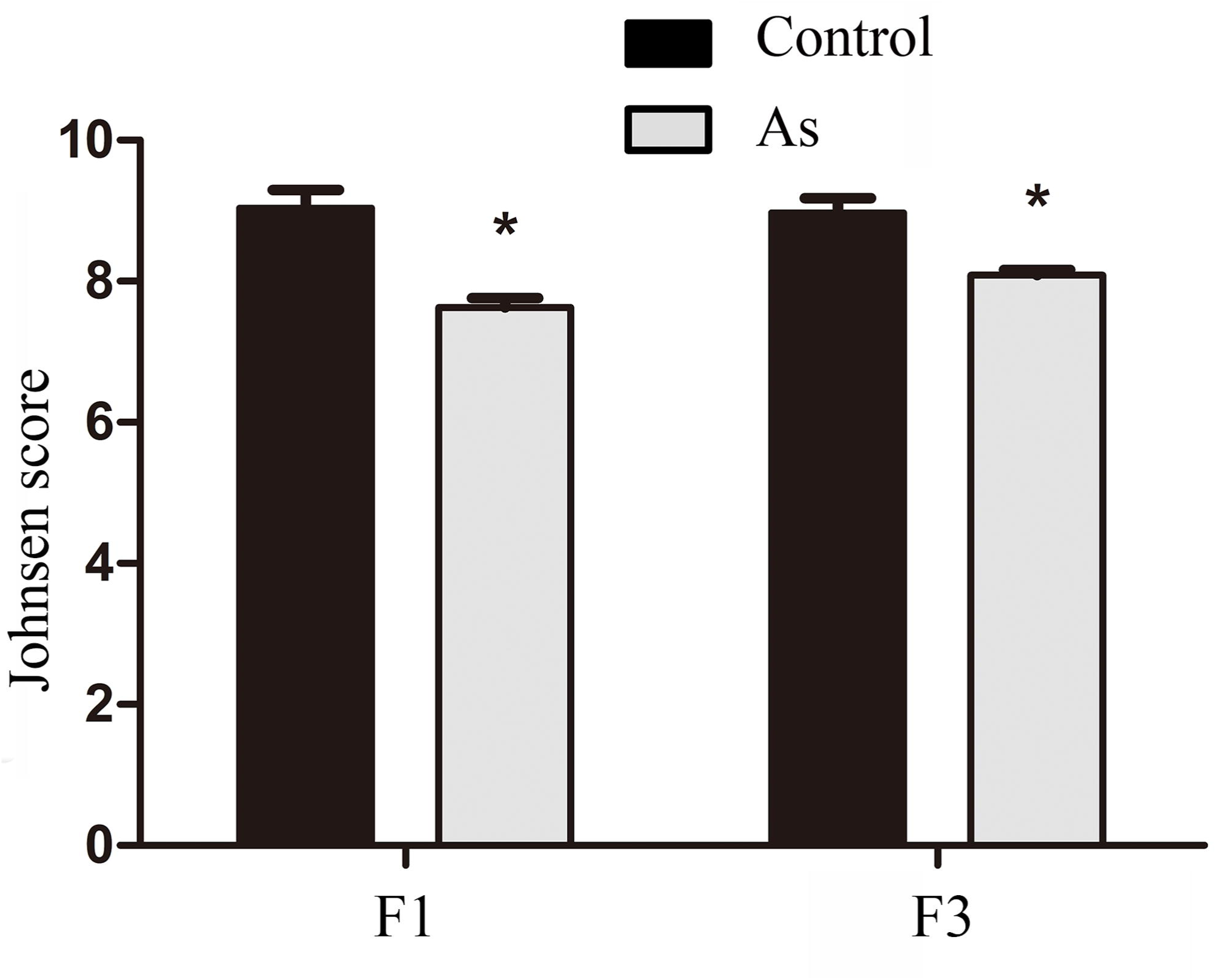
Johnsen score of the arsenic-exposed and control groups in F1 and F3 generations showed difference. Each bar represents the mean ± SEM (*n* = 6). * *p* < 0.05

### 3.5 Epimutations of imprinted genes *Igf2* and *H19* in testis

The methylation status of 14 CpGs of the *Igf2* DMR2 and 15 CpGs of the *H19* DMR was determined by cloning and Sanger sequencing of bisulfite-treated DNA. In the F1 generation, the mean methylation level at the *Igf2* DMR2 was significantly reduced in arsenic-exposed groups from 96.4% to 86.7%. In the F3 generation, the mean methylation level was significantly reduced from 95.2% to 88.8% **(Fig. 5A)**. For the *H19* DMR, the overall methylation level was significantly lower in the arsenic-exposed group than that of the control group in both F1 (96.2% control; 85.3% As) and F3 (96.7% control; 90.9% As) generations (**Fig. 5B**). Individual CpG analysis indicated that arsenic induced hypomethylation of *Igf2* DMR2 and *H19* DMR at several sites in both F1 and F3 generations, but the difference was not significant (**Fig. S1**).

**Fig. 5.**
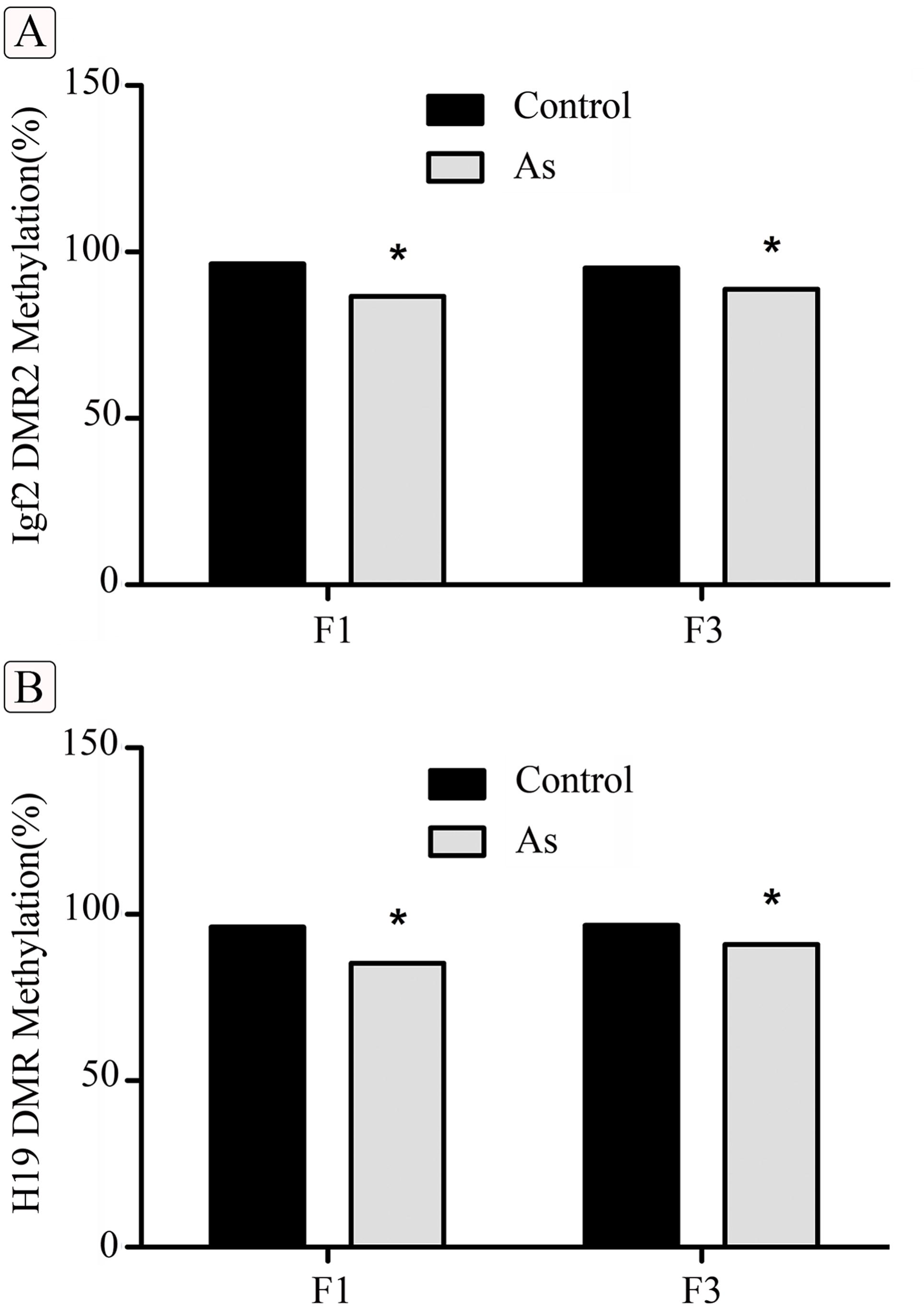
The overall methylation status of *Igf2* DMR2 and *H19* DMR of arsenic-exposed and control groups in F1 and F3 generations. **A** and **B** Statistical analysis of *Igf2* DMR2 and *H19* DMR methylation of arsenic-exposed and control group in F1 and F3 generations (*n* = 3). * *p* < 0.05

### 3.6 mRNA expression of imprinted genes *Igf2* and *H19* in testis

The relative mRNA levels of *Igf2* in testes of arsenic-exposed male offspring in F1 and F3 generations were both significantly higher than those of the control mice (**Fig. 6A**). Showing the same tendency as that of *Igf2* mRNA expression, the relative mRNA expression levels of *H19* in arsenic-exposed males were also significantly higher than those of the control males in both F1 and F3 generations (**Fig. 6B**).

**Fig. 6.**
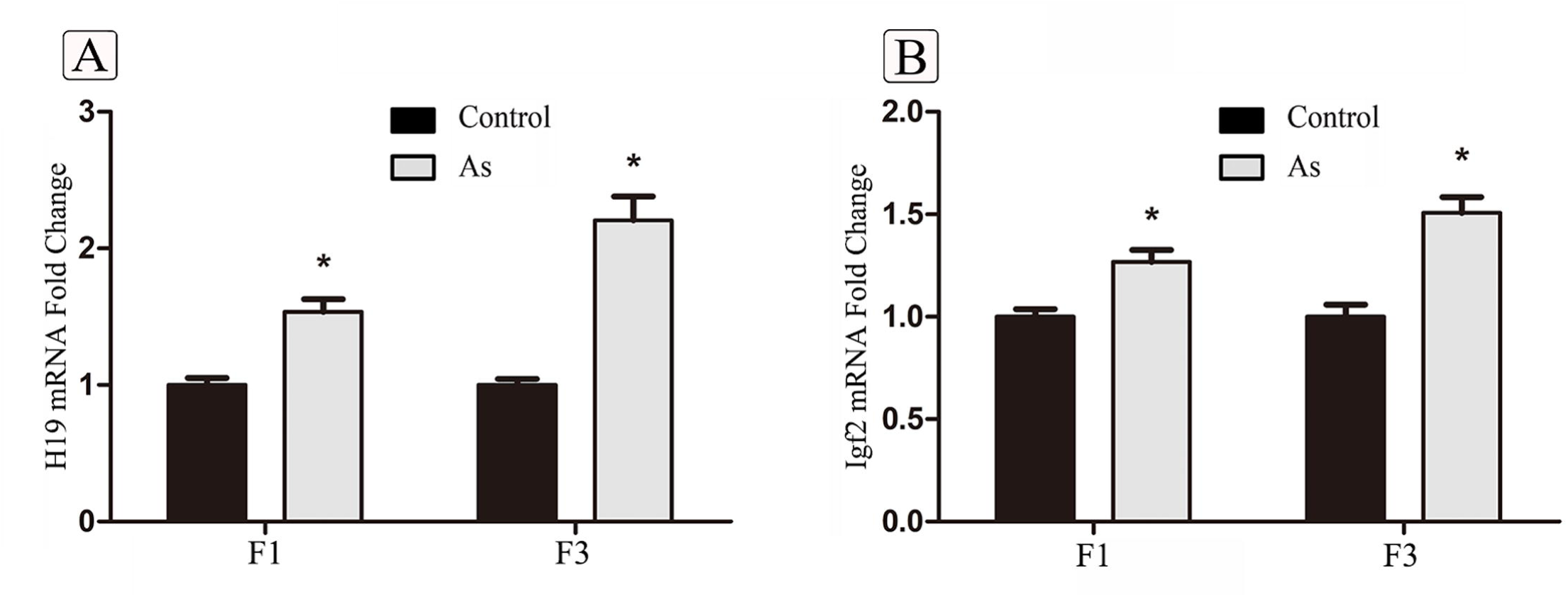
The mRNA expression of imprinted genes *Igf2* and *H19* in testes of male offspring in F1 and F3 generations. **A** Real-time PCR and quantification of *Igf2* mRNA expression levels, **B** Real-time PCR and quantification of *H19* mRNA expression levels. Each bar represents the mean ± SEM (*n* = 6). * *p* < 0.05

## 4. Discussion

In this study, we investigated the transgenerational epigenetic inheritance of *in utero* arsenic exposure (a tumor-inducing 85 ppm level) on reproductive toxicity of male offspring when they reached adulthood. We found that arsenic exposure could cause spermatotoxicity in male offspring of subsequent generations, including reduction of sperm quality and impairment of spermatogenesis, with the epigenetic alteration of imprinted genes *Igf2* and *H19*.

Pregnancy is a particularly sensitive window of susceptibility to environmental insults. This period is critical to germ cell development and epigenetic reprogramming (Ly et al. 2015), and is therefore prone to changes of the *in utero* environment. Exposures that occur during this time can result in permanent and lifelong health effects and can often be passed down to future generations (Hanson and Skinner 2016). Previous studies in CD-1 mice have demonstrated increased tumor incidence after *in utero* exposure to inorganic arsenic at concentration of 85 ppm from GD 8 to 18 (Waalkes et al. 2007). In this study, we focused specifically on the developmental window of susceptibility to arsenic.

For the control pregnant mice (F0), daily water consumption increased during the second-half of gestation. This may be due to the high fetal growth rate during late mouse pregnancy. However, the daily water consumption of arsenic-exposed mice was decreased after the inclusion of sodium arsenite in the drinking water. Compared to the control mice, our data suggested that daily water consumption of the arsenic-exposed mice was reduced by nearly 50% after GD8 (4.73 vs 9.18 mL/day). This result is consistent with a previous report that 50 ppm and 100 ppm sodium arsenite was associated with 47% and 66% reduction in water consumption, respectively (Blakley 1987). The decrease in water consumption may have been related to the decreased body weight gain of arsenic exposed mice **(Fig. 1).**

Since the first observation by Anway et al (2005) that vinclozolin can induce male infertility, a large number of endocrine disruptors have been shown to promote transgenerational inheritance of reproductive disease in laboratory rodents (Manikkam et al. 2013; Zhou et al. 2017). Arsenic is also an endocrine disruptor and male reproductive toxic effects following direct arsenic exposure have been documented in numerous studies. It is generally characterized by a decrease in the testes and accessory sex organ weights and a reduction in sperm count, viability, and motility, as well as degeneration of testicular germ cells (Kim and Kim 2015). Tokar et al. (2012) reported that maternal consumption of methylarsonous acid during pregnancy in CD1 mice produced some unusual testicular lesions in male offspring. In this study, we focused specifically on the first generation of direct *in utero* arsenic exposure (F1 generation) and the first generation of no direct exposure (F3 generation). In line with the aforementioned findings, we found that *in utero* arsenic exposure induced spermatotoxicity by decreasing the sperm motility and increasing the sperm malformation in F1 and F3 generations. Although the testes and epididymides weights between the arsenic-exposed group and the control group were similar, the testes/body weight ratio was decreased in male offspring of arsenic-treated mice both in F1 and F3 generations. The Johnsen scoring system is widely used to evaluate spermatogenesis in the seminiferous tubules (Naeimi et al. 2017). In the current study, reduced Johnsen scores and histopathological alterations were observed in the arsenic-exposed group in both F1 and F3 generations, which indicates poor spermatogenesis. Li et. al (2012) reported that subchronic arsenic exposure has detrimental effects on spermatogenesis and sperm development, which is characterized by a significant reduction in the number of germ cells at stage VII of the spermatogenic cycle in male mice. In this study, a reduction in the numbers of Asg, mPSc, and 7Sd was observed after *in utero* arsenic treatment when compared to the controls. Our data suggest that a transient embryonic exposure to arsenic can cause impairment of spermatogenesis in the F1 generation and part of these phenotypes can be transferred to the F3 generation, indicating a transgenerational inheritance of impaired spermatogenesis. Coincidentally, a more recent study of *C. elegans* showed that the brood size of offspring generations (F1-F5) was significantly reduced by maternal arsenite exposure of the F0 generation (Yu and Liao 2016). Although these two studies were performed in different species, the similar results indicate that maternal arsenic exposure may cause transgenerational reproductive toxicity in subsequent generations.

More observations suggest that ancestral exposure to environmental insults promotes transgenerational inheritance of reproductive disease susceptibility (Nilsson and Skinner 2015). However, the underlying mechanisms remain largely unknown. Recent studies suggest that epigenetic regulations, such as DNA methylation, histone modifications, and small noncoding RNAs, are key mechanisms (Blake and Watson 2016). In an exposed pregnant woman, *in utero* effects on germline epigenetic reprogramming can alter the heritable epigenetic information, resulting in an abnormal phenotype in the offspring. In some cases, these epigenetic changes and linked phenotype can be transmitted to future generations via the germline. Therefore, tracking inherited epigenetic information and its effects for multiple generations is helpful to understand the mechanism of transgenerational transmission of environmentally induced abnormal phenotypes in offspring.

Prenatal arsenic exposure is believed to alter epigenetic programming *in utero* (Cardenas et al. 2015) and the combined *in utero* folate and iAs exposure affected the methylation patterns of CpG islands present in DMRs of imprinted genes (Tsang et al. 2012). Aberrant DNA methylation at imprinted genes, such as *Igf2*/ *H19* and *Mest*, has been associated with male factor infertility in humans (Boissonnas et al. 2010; Santi et al. 2017). In this study, we found a significant decrease in DNA methylation at the *H19* DMR in testes of arsenic-exposed males compared with control males. Arsenic induced a 10.9% and 5.8% decrease in the percentage of methylated CpGs of *H19* DMR in the F1 and F3 male offspring, respectively. The association between male infertility and hypomethylation of *H19* DMR has been well studied. It is reported that the hypomethylation of *H19* may contribute to the decreased spermatogenesis in the offspring after prenatal exposure to vinclozolin and ethanol (Stouder and Paoloni-Giacobino 2010; Stouder et al. 2011). In addition to decreased methylation of *H19* DMR, a significant hypomethylation at *Igf2* DMR2 was also observed in F1 and F3 generations after transient treatment of pregnant CD-1 mice with arsenic. This is in line with other findings of a previous study that transgenerational impaired male fertility was associated with decreased methylation of *Igf2* DMR2 in rat after p,p’-DDE exposure (Song et al. 2014). The impaired methylation of *H19* DMR and *Igf2* DMR2 in testes of F1 adult males may result from the disturbed methylation reprogramming within the developing embryo, because both DNA methylation and arsenic metabolism require S-adenosylmethionine as the methyl donor (Reichard and Puga 2010). This competitive demand for S-adenosylmethionine may affect the percentage of methylated CpG dinucleotides throughout the genome, leading to a change in global and gene-specific DNA methylation. For F3 generation, the altered DNA methylation in *H19* DMR and *Igf2* DMR2 is retained. This may result from a transmission of epigenetic defect via the male germline.

*H19* and *Igf2* are expressed in a monoallelic fashion from the maternal and paternal chromosomes, respectively. Methylation of *H19* DMR and *Igf2* DMR2 plays a critical role in *Igf2* and *H19* allelic transcription. On the maternal allele, CTCF insulator binds to the unmethylated *H19* DMR and blocks the access of enhancer to the *Igf2* promoter, resulting in *H19* expression and *Igf2* repression. On the paternal allele, *H19* DMR is methylated and binding of CTCF is blocked, thus inactivating *H19* and promoting *Igf2* expression (Doshi et al. 2013). DMR2 is an *Igf2* enhancer located in the sixth exon and a deletion of DMR2 could reduce *Igf2* mRNA expression level (Murrell et al. 2001). In our study, hypomethylation of *H19* DMR in testis occurs among arsenic-treated lineage mice, thus resulting in *H19* overexpression. Although both *H19* DMR and *Igf2* DMR2 are hypomethylated, the mRNA expression levels of *Igf2* in the testis are increased in arsenic-treated lineage. In addition to *H19* DMR and *Igf2* DMR2, *Igf2* DMR0 and *Igf2* DMR1 are also involved in the regulation of *Igf2* expression. Constancia et al. (2000) reported that deletion of DMR1 results in biallelic expression of *Igf2*. A recent study indicates that DMR0 is methylated on the active paternal allele in all tissues and may function similarly to mouse DMR1(Murrell et al. 2008). It has been reported that exposure of mouse preimplantation embryos to 2,3,7,8-tetrachlorodibenzo-pdioxin, or TCDD, increased the methylation of *Igf2*/*H19* imprint control region and decreased the mRNA expression of imprinted genes *Igf2* and *H19* (Wu et al. 2004). The heterogeneity of *Igf2* expression may be caused by different environmental exposures and further studies are needed to clarify the interaction between methylation/expression defects of *H19* and *Igf2* of *in utero* arsenic exposure.

## 5. Conclusions

The findings of the present study suggest that abnormal methylation of *H19* DMR and *Igf2* DMR2 and changes in their mRNA expression in testis may play a role in transgenerational impairment of spermatogenesis after *in utero* arsenic exposure. To our knowledge, this is the first study of transgenerational male reproductive toxicity of prenatal arsenic exposure in rodents to date. Considering the differences between rodents and humans, our findings require confirmation but are worthy of further exploration, given their potential implications for the long-term health of arsenic-exposed mothers and their future generations.

## Acknowledgments

This work was supported by National Natural Science Foundation of China [grant numbers 81302390]; Natural Science Foundation of Tianjin City [grant numbers 18JCYBJC91900]; and China Postdoctoral Science Foundation [grant numbers 2016T90210].

