## Supplementary material for "Transgenerational inheritance of abnormal spermatogenesis with *Igf2/H19* epigenetic alteration in CD1 mouse induced by *in utero* arsenic exposure": Figure S1

**Supplementary figure:**

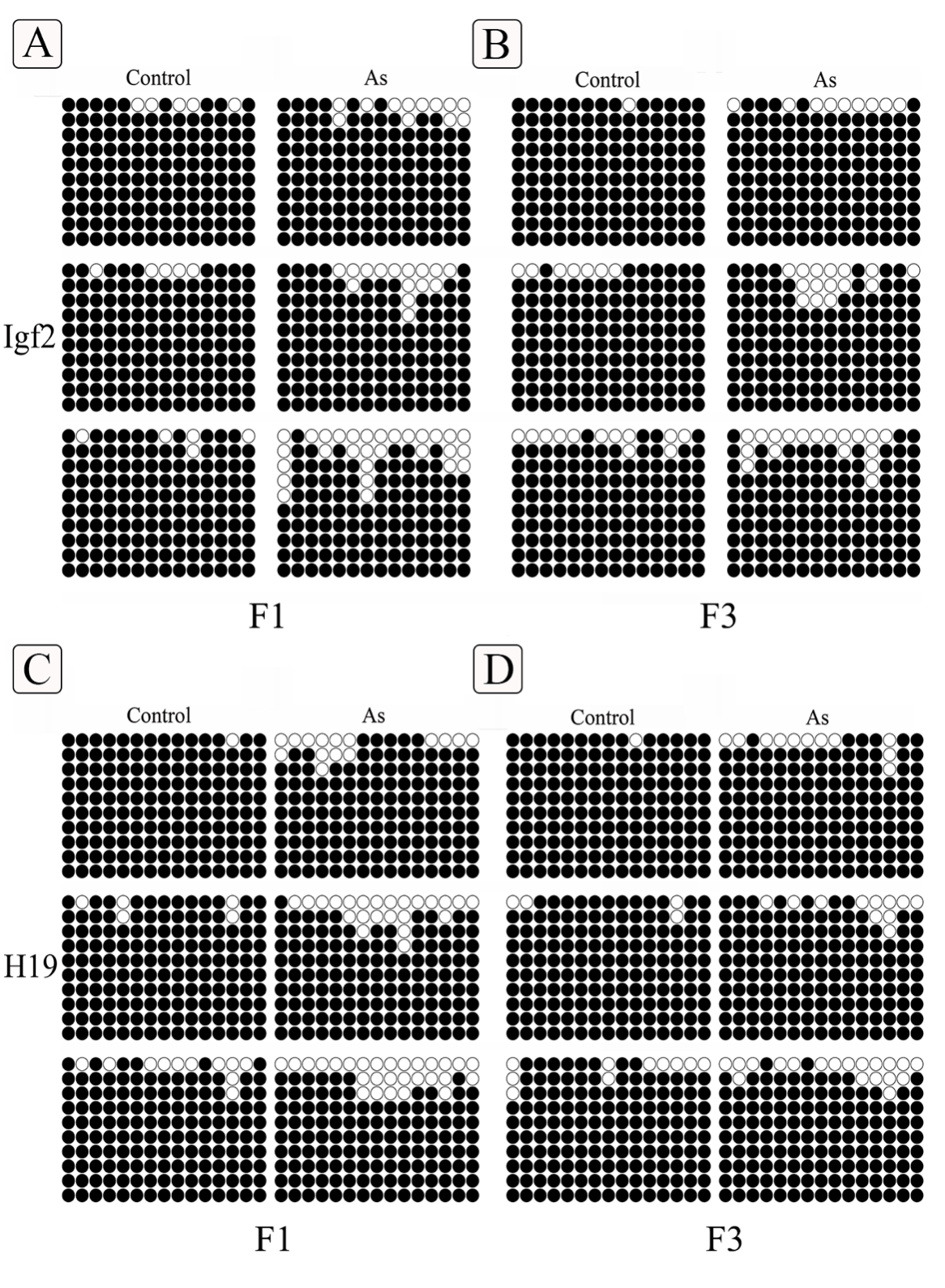

**Fig. S1** The methylation profiles of Igf2 DMR2 and H19 DMR of arsenic-exposed and control groups in F1 and F3 generations. A, B, C, D Methylation profiles assayed by a bisulfte sequencing PCR. Each circle within the row represents a single CpG site (open and closed circles represent unmethylated and methylated CpGs, respectively).
