## Supplementary material for "Transgenerational inheritance of abnormal spermatogenesis with *Igf2/H19* epigenetic alteration in CD1 mouse induced by *in utero* arsenic exposure": Table S1

**Table S1** The criteria of Johnsen scoring system

| Score | Criteria |
| --- | --- |
| 10 | complete spermatogenesis and perfect tubules |
| 9 | many spermatozoa present but disorganised spermatogenesis |
| 8 | only a few spermatozoa present |
| 7 | no spermatozoa but many spermatids present |
| 6 | only a few spermatids present |
| 5 | no spermatozoa or spermatids present but many spermatocytes present |
| 4 | only a few spermatocytes present |
| 3 | only spermatogonia present |
| 2 | no germ cells present |
| 1 | neither germ cells nor Sertoli cells present |
