## Supplementary material for "Transgenerational inheritance of abnormal spermatogenesis with *Igf2/H19* epigenetic alteration in CD1 mouse induced by *in utero* arsenic exposure": Table S2

**Table S2** The litter size and female/male ratio of F1 and F3 offspring after ancestral arsenic exposure (*n* = 6, mean ± SD)

| Generation | Group | Litter number | Litter size | Female/male ratio |
| --- | --- | --- | --- | --- |
| F1 | Control | 6 | 13.17 ± 1.33 | 1.18 ± 0.58 |
|  | As | 6 | 12.00 ± 2.61 | 1.89 ± 1.02 |
| F3 | Control | 6 | 13.00 ± 1.67 | 1.20 ± 0.52 |
|  | As | 6 | 11.67 ± 0.82 | 1.08 ± 0.53 |
