## Supplementary material for "Transgenerational inheritance of abnormal spermatogenesis with *Igf2/H19* epigenetic alteration in CD1 mouse induced by *in utero* arsenic exposure": Table S3

**Table S3** Weight and organ coefficient of testes and epididymides in F1 and F3 generations (*n* = 15-18, mean ± SD)

| Generation | Group | Epididymides  (g) | Epididymides/  body weight (%) | Testes  (g) | Testes/  body weight (%) |
| --- | --- | --- | --- | --- | --- |
| F1 | Control | 0.14 ± 0.03 | 0.46 ± 0.09 | 0.27 ± 0.03 | 0.87 ± 0.14 |
|  | As | 0.16 ± 0.03 | 0.41 ± 0.07 | 0.29 ± 0.04 | 0.75 ± 0.09* |
| F3 | Control | 0.15 ± 0.05 | 0.40 ± 0.14 | 0.27 ± 0.03 | 0.72 ± 0.09 |
|  | As | 0.16 ± 0.03 | 0.44 ± 0.10 | 0.25 ± 0.05 | 0.68 ± 0.13 |

*Compared with the control group: *p* < 0.05
